# Decipher: A computational pipeline to extract context-specific mechanistic insights from single-cell profiles

**DOI:** 10.1101/2024.05.01.591681

**Authors:** Edgar Basto, Bilal Wajid, James Read, Jesse Armitage, Jason Waithman, Michael Small, Anthony Bosco

## Abstract

The advent of single-cell profiling technologies has revolutionized our understanding of the cellular and molecular states that underpin disease. However, current computational tools struggle to recover both known and novel mechanistic insights at distinct layers of biological regulation. Here, we present *Decipher*, a novel computational pipeline that builds integrated cell signalling networks from single-cell profiles in a context-specific, data-driven manner and identifies the key cellular and molecular events that drive disease. We benchmarked *Decipher* against existing tools and found it could recover known, experimentally determined cytokine signalling pathways, whilst maintaining the flexibility to detect novel pathways and context-specific effects. Notably, *Decipher* produces global cell-to-cell signalling maps that are interpretable. We utilised *Decipher* to unveil the cellular and molecular mechanisms driving a novel population of inflammatory monocytes enriched with interferon stimulated genes that is markedly increased in frequency following secondary immunization with the Pfizer-BioNTech COVID-19 mRNA vaccine. Finally, we employed Decipher to interrogate regulon profiles from covid-19 patients with mild versus severe disease, and we found that progression to severe disease was associated with a loss of interferon signalling transcription factors (Irf7, Irf9, STAT1, STAT2) and a gain of factors that drive inflammation and cellular stress responses (NFkB, HIF-1a, ATF3, ATF4). Taken together, our findings demonstrate that *Decipher* can decode signalling pathways and report on ligand-receptor mediated transcription factor-target gene networks that underlie processes in homeostasis, disease, and cellular responses to therapies. We present *Decipher* as an invaluable new tool for the discovery of novel therapeutic targets and the development of new medicines.

## 1 Main

Complex multicellular life forms execute a multitude of high-level biological functions, including growth, differentiation, metabolism and homeostasis. These functions are mediated by interactions among multiple cell types through diverse molecules, such as ligands and receptors^1^. The advent of single-cell omics, which profiles biological systems at single-cell resolution^2^, along with computational tools that infer patterns of cell-to-cell communication from such data^3,4^, have furthered our understanding of the role of intercellular interactions in diverse contexts such as the maternal-fetal interface^5^, wound healing^4^, human development^6^ and cancer^7^.

While inferring cell-cell communication patterns is an active area of method development, with over one-hundred available tools catalogued by Armingol et al.^8^, most share two key limitations: they overlook the downstream transcriptional effects of upstream signalling events^9^, and even those that consider such effects often rely heavily on pre-defined knowledge graphs to connect upstream signalling to cellular responses.

Accounting for downstream effects is essential, as it reveals how cellular interactions reprogram gene expression and ultimately drive disease processes^7^, insights which are lost when analyses stop at ligand-receptor pairs alone. Conversely, integrating curated knowledge graphs can improve the reliability and accuracy of inferred networks^10^, but at the same time, they can also bias the results towards well-studied pathways^11^, and obscure novel or context-specific signalling.

We surveyed eleven computational tools that analyse cell-cell communication and explicitly account for downstream signalling. We found that most methods tend to rely on curated molecular interaction databases, limiting their findings to well-studied relationships. This is the case for Pathway-centric methods (SoptSC^12^, CommPath^13^), Multilayer knowledge-graph methods (NicheNet^7^, LRLoop^14^, Scriabin^15^) and Transcription-factor-centric methods (ScMLNet^16^, CellCall^17^, scSeqComm^18^, SPARTAN^19^). *De novo* network inference tools (CytoTalk^20^, DIALOGUE^21^), in contrast, avoid this reliance on prior knowledge by reconstructing networks directly from expression data, yet the resulting relationships often lack mechanistic context. In both cases, the methodology limits the ability to identify novel signalling pathways.

To address these gaps, we developed *Decipher*, a novel computational platform that builds integrated signalling networks that operate between and within cells, capturing both ligand-receptor signalling and transcription factor-target gene regulatory activity. *Decipher* utilizes prior-knowledge approaches when considering these networks separately, and data-driven network inference when integrating them. Thus, *Decipher* carefully balances the ability to accurately prioritize signalling mechanisms with the necessity to detect previously unidentified pathways.

## 2 Results

*Decipher* requires annotated single-cell RNA-seq profiles from two experimental conditions, a database of interacting ligand-receptor (LR) pairs and a reference gene regulatory network capturing relationships between transcription factors (TF) and their downstream target genes. A *Decipher* analysis is not limited to any specific source of prior knowledge, however, by default we use LR pairs from connectomeDB2020^22^ and a base network from CellOracle^23^. In addition, as gene regulatory network wiring is context-specific^24^, we leverage further functionality from CellOracle^23^ to tailor the base network to the experimental and cellular context of each cell type, retaining only the edges that show evidence of being active.

To score LR signalling, Decipher relies on reconstructed integrated signalling networks to link intercellular signalling potential to downstream transcription factor activity. Signalling potential refers to the capacity of cells to engage in differential signalling and is estimated as the mean-product^22^ of ligand and receptor expression across meta-cells^25^ (a strategy we employ to limit dropout effects). Transcription factor activity reflects the influence TFs exert within the cell and is inferred by applying an overdispersion-based scoring function from PAGODA2^26,27^ to the expression profile of the target gene set for each transcription factor, as per the tailored gene regulatory networks. To reconstruct integrated signalling networks, the calculated interaction potentials and TF activities are used as the predictor and target variables in a random forest regression model, whose feature importance values represent the edge weights connecting LR pairs to TFs within the network. The sign of the Spearman correlation between each predictor-target pair designates the mode of influence of that edge. Together, these steps yield a weighted, signed integrated signalling network that captures how extracellular cues propagate to transcriptional responses.

To quantify the activity of each LR pair *i* across conditions, *Decipher* integrates the predicted regulatory weight *w*_*i,f*_ between LR pair *i* and transcription factor *f* with the observed changes in TF activity Δ_*f*_, defined as the difference in median TF activity between conditions. The resulting *Decipher* score *S* is computed as:

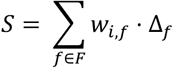

Where *F* denotes the set of TFs relevant to a given cell type. This formulation prioritizes LR pairs that are both influential (high *w_i,f_* indicating a strong regulatory relationship) and associated with responsive TFs (a large Δ_*f*_ signals a substantial intracellular change). *Decipher* scores support three core layers of analysis: cell-to-cell LR signalling, LR-to-TF regulatory signalling and TF-target gene regulation (Fig. 1).

**Figure 1.**
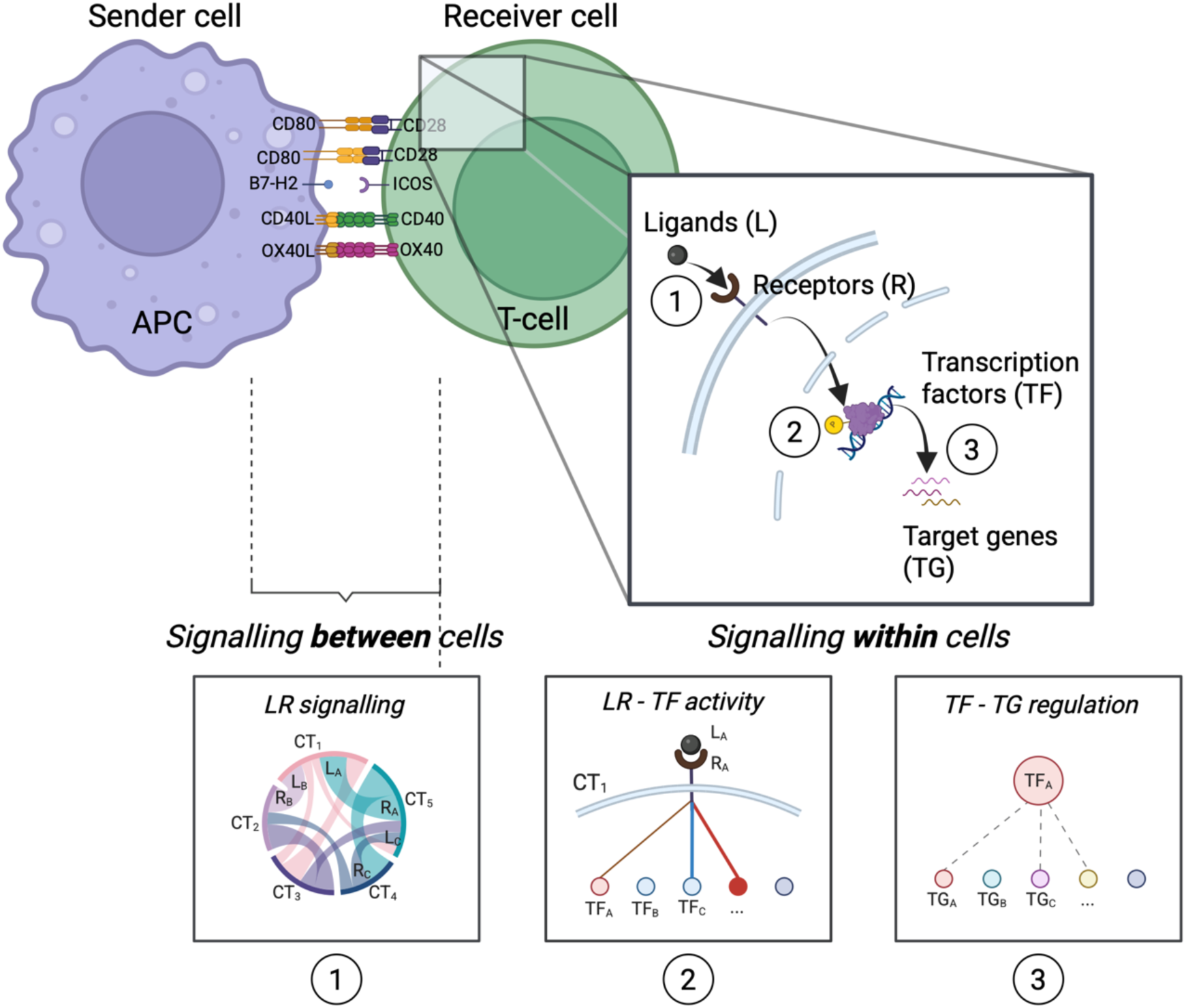
Overview of the *Decipher* framework for integrating intercellular and intracellular signalling. *Decipher* connects ligand-receptor (LR) interactions between a sender cell (e.g., antigen-presenting cell, APC) and a receiver cell (e.g., T-cell) to downstream transcription factor (TF) activity and target gene (TG) regulation. The framework consists of three hierarchical layers: (1) ligand-receptor interactions, (2) the interaction between LR pairs and TF activity profiles, (3) TF-target gene regulation. Together, these layers capture key signalling pathways both between and within cells.

### Benchmarking Decipher with other cell-cell communication methods

To demonstrate the utility of *Decipher* in extracting insights into the cellular and molecular responses to a perturbation, we applied *Decipher* to twelve publicly available scRNA-seq datasets (Methods) focused on immune responses to viral and bacterial infections^28–31^, vaccination^32,33^, autoimmune conditions^34,35^, genetic disorders^36^, chronic disease^37^, and cancer^38,39^. We compared *Decipher* with four other cell-cell communication methods: two that account for intracellular responses (NicheNet^7^ and LIANA+^40^) and two that focus solely on intercellular signalling (Connectome^41^ and NATMI^22^).

We first profiled the general characteristics of the LR interaction scores produced by each method, including the total number of reported LR pairs and the corresponding distribution of their scores. Although the number of reported interactions by each method is not strictly a measure of performance, it can be informative about how a method balances sensitivity and specificity.

We observed that *Decipher* reports two orders of magnitude fewer interactions than NicheNet, NATMI, and Connectome, and one order of magnitude fewer interactions than LIANA+ (Fig. 2a). Methods further differed in the characteristics of the distributions of their prioritization scores (Fig. 2b). Only *Decipher*, LIANA+, and NATMI produced negative and positive scores, which indicate inhibition and activation of signalling, respectively. The score distributions produced by the five methods formed four distinct patterns. *Decipher* displayed a sharp peak at zero with long, exponentially decaying tails. NicheNet had a peak at a moderately positive value and much shorter tails. LIANA+ and NATMI displayed bimodal distributions, with peaks at moderately negative and moderately positive values, with NATMI exhibiting especially short tails. Lastly, Connectome exhibited a sharp peak at zero and a long right tail, though the decay was less steep.

**Figure 2.**
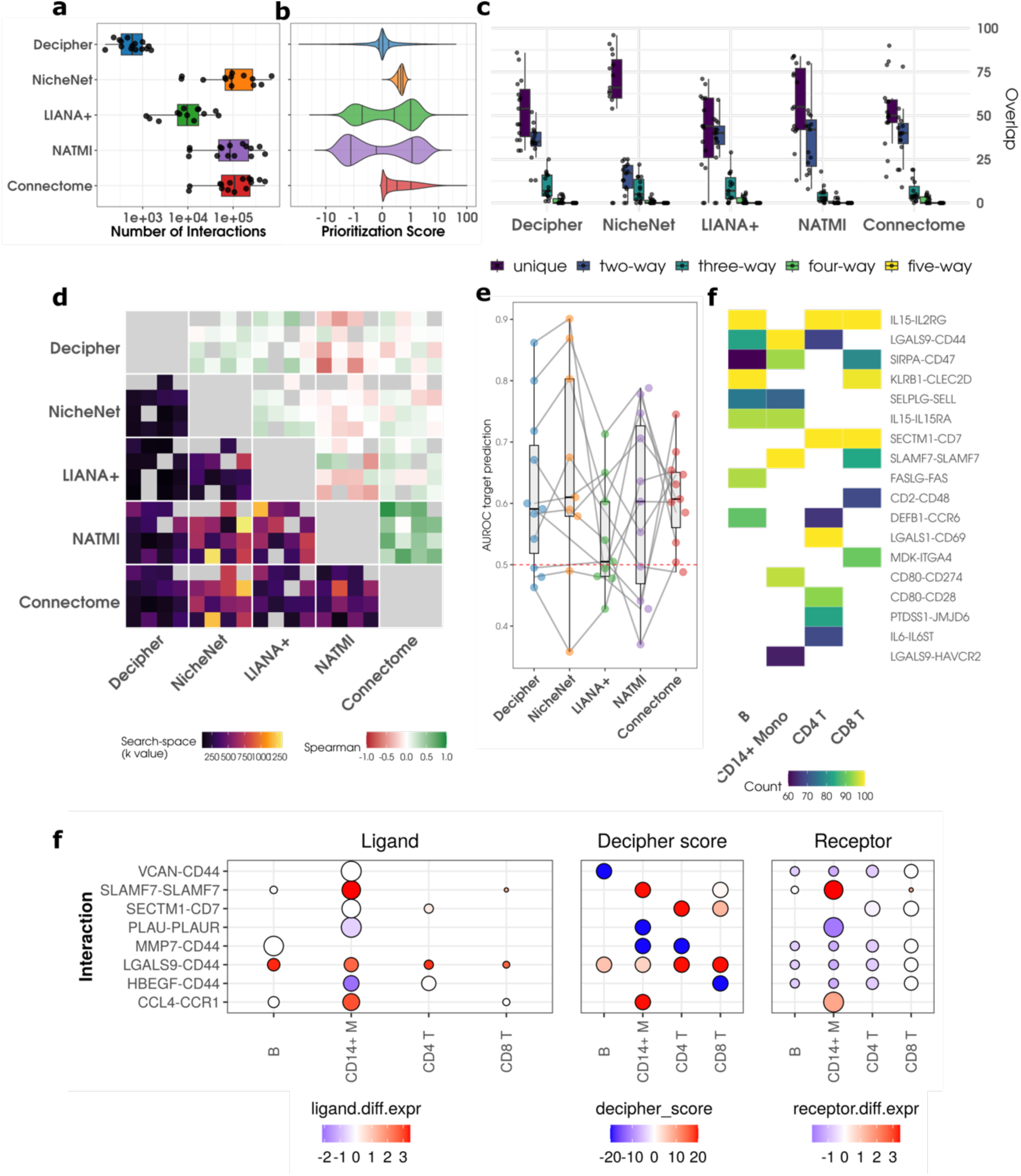
Comparison of predicted ligand-receptor activity across methods. (a) Box plots summarizing the number of reported interactions for each method across all datasets. (b) Violin plots of the combined distributions of predicted interaction scores for each method across all datasets. (c) Box plot of the overlap between methods for the top 100 interactions in each dataset. The plot displays the number of interactions unique to each method, as well as the number of two-way, three-way, four-way, and five-way overlap. (d) Compound heatmap of the search space (bottom-left of heatmap) required to find 100 overlapping LR pairs that are both highly ranked, as well as Spearman correlations (top-right of heatmap) between the rankings of the top 100 overlapping LR pairs. The search space is displayed using a Viridis scale, where darker colours indicate a smaller search space and brighter colours a larger one. A smaller search space indicates a greater degree of agreement between methods. Spearman correlation utilizes a red to green colour scale, where red indicates negative Spearman correlation, green positive correlation, and white no correlation. (e) Beeswarm plot of AUC scores for eleven datasets benchmarking the performance of Decipher and other frameworks against CytoSig scores as true labels. We omitted the lupus and sepsis datasets from this comparison, as CytoSig did not define enough active ligands to construct reliable ROC curves. (f) Heatmap indicating the consistency of interactions prioritized by Decipher across 100 runs on the dataset of lupus patient response to interferon-β^44^ with distinct random seeds. White indicates that a ligand-receptor pair was not relevant to a particular cell type. (g) System-level signalling for the same dataset. The left sub-plot displays ligand-level statistics for the sender cell type, while the right sub-plot presents receptor-statistics for the receiver cell type. The size of the bubbles indicates the relative expression of the gene in the corresponding cell type, whereas the colour indicates the differential expression of that gene in case vs control. The central sub-plot presents *Decipher* scores for all receiver cell types. Here, blue indicates that a LR pair is predicted to be inhibited, and red activated.

Consensus among methods was limited. Overlap between the top-100 ranked LR pairs per dataset (Fig. 2c) showed that NicheNet reported the greatest number of unique interactions. In comparison to other methods, NicheNet displayed few two-way intersections, whereas three-way overlap was most frequent among *Decipher*, NicheNet, and LIANA+. Four- and five-way overlaps were rare. Agreement in ranking was assessed on the top-100 ranked consensus LR pairs using Spearman correlation and search depth (Fig. 2d), defined as the rank position within each method required to retrieve all 100 consensus interactions. *Decipher* required the smallest search depth in every pairwise comparison. LIANA+ displayed a similar profile to *Decipher*, except when compared with NATMI. NATMI and Connectome showed moderate mutual agreement, while NicheNet required markedly deeper searches to align with NATMI and Connectome. Spearman correlation further highlighted structural differences among methods. NATMI and Connectome displayed the highest consensus among all pairs. *Decipher*, NicheNet and LIANA+ were weakly to moderately correlated with one another. In contrast, NATMI was negatively correlated with this group.

We benchmarked each method against CytoSig^42^, which predicts cytokine activity for 62 ligands based on experimentally determined gene expression signatures. Because NicheNet shares training data with CytoSig^43^, we treated NicheNet as a positive control. In some datasets, CytoSig detected an insufficient number of active ligands (median z-score ≥ 2) to construct reliable ROC curves. *Decipher*, NicheNet and NATMI achieved the highest upper-range AUC values, although the latter two exhibited a few low-performing outliers. Connectome showed intermediate performance with modest variance, whereas LIANA+ displayed limited predictive power in this benchmark.

Because *Decipher* includes stochastic steps, we tested its robustness to changes in the initial random seed. The analysis was repeated 100 times on a dataset of lupus patient response to interferon-β^44^ distributed through the ExperimentHub package from Bioconductor^45^. Across the majority of the 100 runs, the same LR pairs were consistently prioritized (Fig. 2f); for example, IL15-IL2RG dominated B, CD4 T, and CD8 T cell rankings, whereas LGALS9-CD44 was frequently the leading interaction for CD14^+^ monocytes.

Lastly, we compared the visual outputs of *Decipher*, NicheNet and LIANA+. Each tool conveys two metrics: the signalling potential of individual ligands and receptors, and an activity score for every LR pair. To convey the signalling potential, *Decipher* reports on the normalized proportion of transcripts originating from each cell type, whereas NicheNet relies on the average expression of each ligand and receptor. LIANA+ does not convey this information. Differential signalling, on the other hand, is captured by all three methods by reporting on the differential expression of the ligand or receptor separately. Because *Decipher* aggregates all sender cell types into a single mixed cluster, it presents fewer visual outputs than the other methods. Reformatting NicheNet and LIANA+ outputs for the infant response to Poly-IC comparison to the same visual output format as *Decipher* highlighted this increase in complexity (Supplementary Fig. SF.1).

### Extracting mechanistic insights from predictions made by *Decipher*

To assess the capacity of *Decipher* to extract mechanistic insights and identify candidate therapeutic targets from single-cell profiles, we focused on two studies related to COVID-19: a study investigating the immune response to the Pfizer-BioNTech COVID-19 vaccine in humans^33^ and a study profiling PBMCs from patients with mild or severe COVID-19^31^. Both studies included baseline, unvaccinated or healthy, controls.

In the vaccination study, Arunachalam et al.^33^ described the strong activation of the innate immune system and the emergence of a novel CD14^+^BDCA1^+^PD-L1 (cluster C8) monocyte subpopulation one day post-secondary vaccination. Moreover, Arunachalam et al. characterized the intracellular activity of this subpopulation, identifying activation of STAT1, STAT2, STAT3, IRF1 and IRF8 TFs, and associatively linked the observed response to interferon gamma (IFNG) signalling, hypothesizing that these two factors were a key component of the programmed immune response after vaccination.

We applied *Decipher* to identify the upstream and downstream signals driving activation of the C8 monocyte subpopulation. The intracellular response of the C8 cluster showed strong upregulation in subsets of the IRF and STAT family of TFs: IRF2, IRF7, IRF8, STAT1, and STAT2, consistent with observations reported by Arunachalam et al.^33^, (Fig. 3a). *Decipher* also detected IFNG signalling into the C8 subpopulation (Fig. 3b), in further accordance with the reference study^33^. In addition, we observed autocrine and paracrine signalling into the C8 monocytes through SLAMF7, CCL2, and C1QB pathways, as well as inhibited signalling via PRG4-TLR2 and HGF-CD44 pathways (Fig. 3b). We further characterized the downstream TF response to prioritized LR pairs and observed that C1QB, CCL2, and SLAMF7 activation are strongly related to STAT2, IRF2, IRF7, and IRF8 activity, as well as moderately related to a much broader transcription factor response (Fig. 3c). IFNG, as well as other activated pathways identified by *Decipher,* such as SPN and HGF, exhibited a more exclusive response.

**Figure 3.**
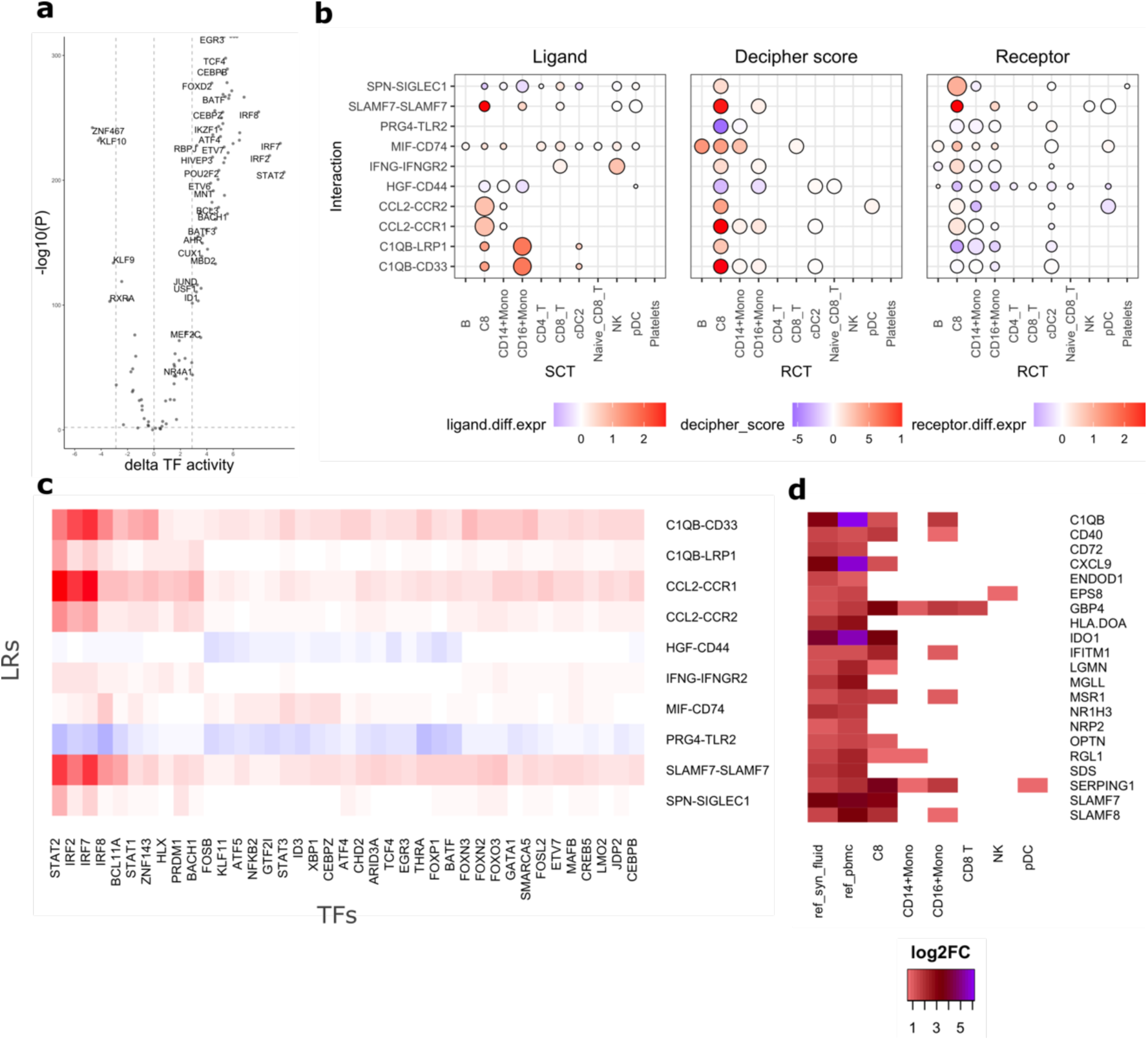
Mechanistic insights in the Pfizer–BioNTech COVID-19 vaccine comparison. (a) Volcano plot of changes in transcription factor (TF) activity in the C8 cluster on day one post-secondary vaccination vs baseline. (b) Multi-panel plot of global dynamics from the top 10 identified ligand–receptor (LR) pairs in the C8 cluster. (c) Heatmap of individual LR–TF *Decipher* scores for the LR pairs highlighted in the *Decipher* multi-panel plot. (d) Heatmap of reference log fold-changes from the SLAMF7-high gene signature in synovial fluid (ref_syn_fluid) and PBMCs (ref_pbmc) from sorted CD14^+^ monocytes (Simmons et al.^46^), shown alongside differential expression profiles from three monocyte populations (C8, CD14^+^, and CD16^+^), as well as CD8 T, NK, and pDC cells.

To validate the predicted SLAMF7 signalling, we compared differential expression signatures of C8 monocytes, CD16^+^ monocytes, CD8 T, NK, and pDC cells with a published signature of SLAMF7^high^ macrophages from synovial fluid and PBMCs in an inflammatory setting^46^. The C8 signature, and to a lesser extent the CD16^+^ signature, matched the reference (Fig. 3d). Although pDCs, NK cells and CD8 T cells express SLAMF7, their differential expression signatures did not match the reference signatures, nor did *Decipher* identify them as receiving signalling via SLAMF7.

We next compared mild and severe COVID-19 samples. Cell types were annotated with Azymuth^47^ using its PBMC reference annotated dataset. Analysis was restricted to cell types that were sufficiently abundant and exhibited either changes in differential expression or in proportion across conditions (Fig. 4a). Out of the 29 cell types identified by Azimuth, fourteen represented less than 1% of the total cell pool and were excluded. Abundance differences were further assessed with scCODA^48^, which identified six cell types that presented significant changes in abundance (FDR < 0.05): B naive cells, CD16^+^ monocytes, CD4 TCMs, CD8 cells, NK cells, and plasmablasts. To elucidate changes in these populations, we calculated the total number of differentially expressed genes (DEGs), as well as the proportion corresponding to mild versus severe and up-vs down-regulation (Fig. 4c).

**Figure 4.**
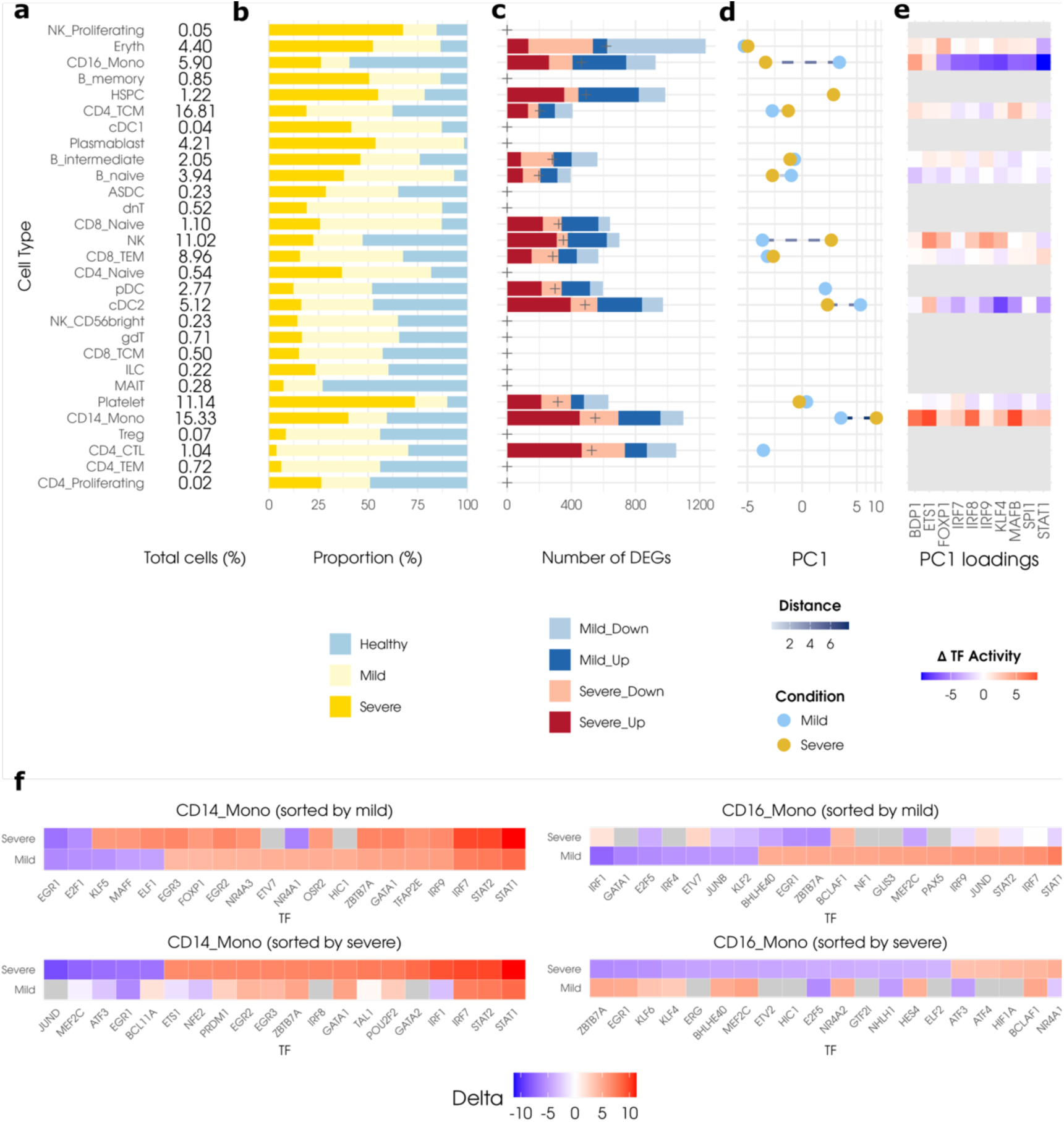
Cell-type-specific transcriptional differences in mild and severe COVID-19. (a) Text summarizing the total proportion of each cell type across the three conditions (Healthy, Mild, and Severe). (b) Bar plot of the relative proportion of each cell type for each condition. Guide vertical lines are displayed at the 33% and 66% values. (c) Bar plot of total number of differentially expressed genes and changes in the proportion of upregulated vs downregulated genes. Crosshairs in each bar indicate the 50% mark of the number of differentially expressed genes. (d) Line plot of the first principal component embedding of changes in TF activity by cell type and condition. Cell types across conditions are joined by a dotted line whose colour reflects the distance (Euclidean) between them. (e) Heatmap of differences in changes in TF activity (mild – severe) for top 10 PC1 loadings. (f) Differential TF activity plots for CD16^+^ Monocytes and CD14^+^ Monocytes, sorted by absolute value of change in TF activity for that given cell type in either the mild or severe condition.

We then quantified changes in transcription factor (TF) activity in the two comparisons and projected the change in TF activity scores onto a reduced-dimensional representation via principal component analysis. Four cell types: CD14^+^ monocytes, CD16^+^ monocytes, NK and CDC2 cells, showed the largest differences in the first principal component embedding between the mild and severe conditions (Fig. 4d). The loadings corresponding to the first principal component accounted for 36.7% of the observed variance and included interferon-related transcription factors IRF7, IRF8, IRF9 and STAT1 (Fig. 4e). Critically, a cell-type- and condition-dependent divergent response to mild and severe COVID-19 was apparent in the direction of change along the PC1 axis. Heatmaps of the delta in the change in TF activity between mild and severe, calculated as the subtraction of the mild scores from the severe scores, further showcased evidence for this divergent response; CD16^+^ monocytes in severe, for example, exhibited inhibition in most TFs related to PC1, whereas CD14^+^ monocytes in severe exhibited strong activation of those same TFs (Fig. 4e). Together, these observations suggest that these cell types underwent substantial transcriptional reprogramming, particularly in interferon-related functions, between the mild and severe states.

To further corroborate this observation, we visualized changes in TF activity sorted by the absolute values for both mild and severe conditions for CD14^+^ and CD16^+^ monocytes (fig. 4f). Here, we observed that CD16^+^ monocytes from the mild condition display strong differential activation of interferon-related transcription factors (STAT1, IRF7, STAT2, JUND, and IRF9), which were not upregulated in CD16^+^ monocytes from the severe condition. This difference appears reversed in CD14^+^ monocytes, albeit the effect does not speak to a divergent response, but rather to a change in intensity of activation.

To further characterize this divergence in intracellular programs we examined comparative network representations across the three layers produced by the *Decipher* analysis for both the mild and severe comparisons of the two monocyte subpopulations (Fig. 5). The LR signalling of these networks showed that CD14^+^ monocytes in the severe condition exhibited the strongest activation of intercellular signalling: via CD63, SIGLEC1, SELL and ITGB2 receptors. In addition, pDCs only played a role in signalling in the mild condition. We again observed markers of type I interferon signalling via the TFs IRF7, IRF9, STAT1 and STAT2, active in CD14^+^ monocytes from both mild and severe COVID, and in CD16^+^ Monocytes from mild COVID only. In CD16^+^ monocytes from severe COVID, there is a complete loss of type I interferon signalling TFs. Moreover, this loss of type I interferon TFs is associated with a gain in TFs that drive inflammation (NFkB2) and responses to oxidative and cellular stress (HIF-1A, ATF3, ATF4).

**Figure 5.**
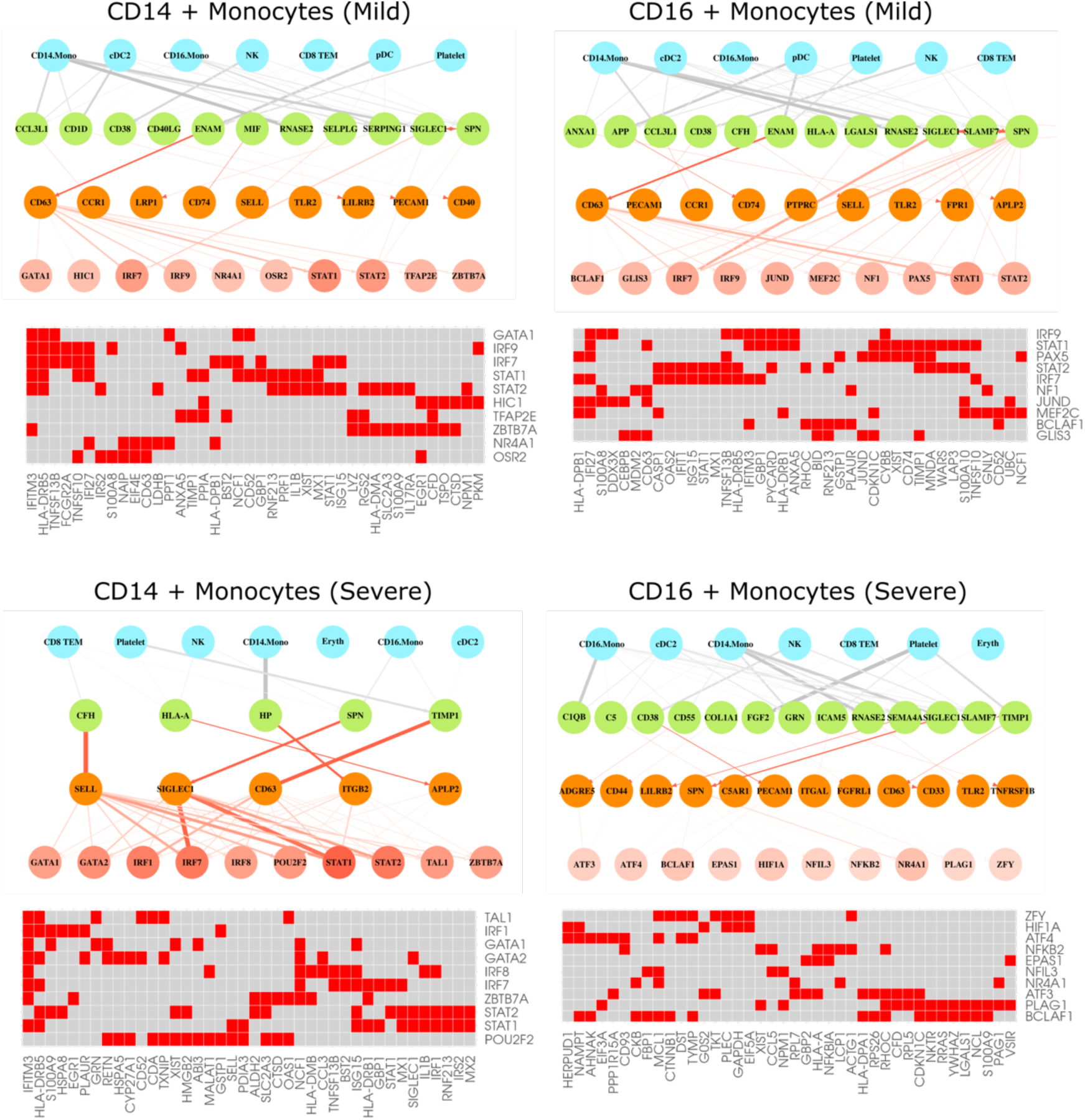
Multi-layer visualizations of *Decipher* results for CD14^+^ Monocytes and CD16^+^ monocytes in mild and severe COVID-19. Network graphs displaying sender cell type (blue), ligand (green), receptor (orange) and transcription factors (TFs) for the CD14^+^ and CD16^+^ monocytes in mild and severe COVID-19. Each network is constrained to the top 10 upregulated TFs, coloured by the change in calculated TF activity. Edges between sender cell types and ligands are weighted by the proportion of total ligand expression (normalized by cell count) expressed by each cell type. Additionally, only the top three cell types producing a given ligand are given an edge with that ligand. In addition, only ligands and receptors involved in the top 20 interactions by Decipher score and strongly associated with the top TFs (based on permutation importance) were considered. Lastly, each network is accompanied by a heatmap of downstream target genes (TG) for each TF. We limited these target genes to the top 40 most researched genes (within the set of target genes for the selected TFs), as determined by the number of publications that mention said gene in PubMed^49^. Red colour within the heatmap indicates the existence of a regulatory relationship between a TF and a TG. All four plots share the same scale for the node colours (applicable to TFs) and widths or colours of edges.

## 3 Discussion

Decoding the cellular and molecular events that drive disease processes and cellular responses to therapies requires the development of new methods that can extract deep mechanistic insights at multiple layers of regulation from single-cell profiles. Cell-cell communication is one of these critical layers, as it mediates many of the functions necessary for multicellular life. However, among the methods that study intercellular communication, most only focus on upstream ligand-receptor interactions and fail to consider downstream effects. Among the subset of methods that do account for multiple regulatory levels, most tend to overly rely on prior knowledge, limiting their findings to well-characterized relationships. Those that do rely primarily on data-driven network reconstruction often lack mechanistic context and may not even possess sufficient predictive power for network inference^50^. In both instances, the methodology limits the ability to identify novel signalling pathways. Toward this goal, we developed *Decipher*, a novel computational pipeline that builds integrated cell-signalling networks from single-cell profiles and unveils key cellular and molecular events that drive biological responses. *Decipher* employs both prior knowledge and data-driven approaches to strike a balance between novelty and accuracy. We found that *Decipher* performed as well as or better than other state-of-the-art cell-cell communication methods, while also generating more interpretable visualizations. To test the ability of *Decipher* to extract mechanistic insights, we analysed two COVID-19 single-cell datasets. In the first study, we identified the upstream signals that drive a previously described monocyte subpopulation critical to the immune response post Pfizer-BioNTech COVID-19 vaccination. In the second study, we characterized the difference in cellular responses of monocytes in mild and severe COVID-19, which aligned with current understanding of the mechanisms of this disease. Taken altogether, our findings suggest that *Decipher* can decode signalling pathways and report on mechanistic relationships captured through ligand-receptor mediated transcription factor – target gene networks.

We benchmarked *Decipher* against four established cell-cell communication methods and found that *Decipher* returns a tightly prioritized set of molecular pathways with scores that capture both the strength and mode of influence across ligands, receptors, transcription factors and target genes. While agreement among methods was generally low, as previously noted for other methods by Dimitrov et al.^11^ we did observe that tools limited to exploring only ligand-receptor interactions behaved differently from those that included intracellular signalling as well. When evaluated against ligand activity predictions from CytoSig^42^, *Decipher* performed as well as or better than other methods, while maintaining the flexibility to detect novel pathways by reducing its reliance on prior knowledge. Importantly, *Decipher* consistently prioritized top ligand-receptor pairs across runs, suggesting robustness in its analysis. We further observed that, by pooling all sender cell types into a single mixed cluster, *Decipher* produced signalling maps that reduce visual clutter without sacrificing biological insight. Thus, *Decipher* offers a balance between highly prioritized interactions, concise visualisations and competitive predictions.

To evaluate the capacity of *Decipher* to extract mechanistic insights from single-cell profiles, we analysed two COVID-19 scRNA-seq studies profiling PBMCs: the first from individuals one day post-secondary Pfizer-BioNTech COVID-19 vaccination, and the second from patients with mild or severe COVID-19. We employed *Decipher* to recover the underlying mechanism driving a novel subpopulation of CD14^+^BDCA1^+^PD-L1^+^ monocytes identified by Arunachalam et al. to be markedly increased in frequency following secondary vaccination and characterized by Type I interferon response, TLR and inflammation, with elevated activity of interferon and STAT TFs and reduced activity of AP-1 TFs^33^. *Decipher* analysis confirmed the previously reported upregulation of IFN (IRF2, IRF7, IRF8) and STAT (STAT1, STAT2, STAT3) family of TFs, as well as displayed significant crosstalk between innate immune cell types, as well as autocrine signalling in C8 cells, a common phenomenon in cell-cell communication^3^. We confirmed previous data demonstrating that interferon gamma (IFNG) is a driver of the C8 population^33^, as well as identified CD8 T and NK cells as the primary producers of this ligand^51^. IFNG, however, was not the strongest signal found by *Decipher*, as we also found significant signalling through the SLAMF7, CCL2 and C1QB ligands into the C8 cluster. Notably, SLAMF7 is a self-ligand receptor from the signalling lymphocytic activation molecule (SLAM) family and a potent regulator of interferon responses^52^ and inflammatory macrophage activation^46^. Importantly, IFNG and SLAMF7 have been previously implicated as a driver of a super-activated macrophage state present in autoimmune diseases and severe COVID-19^46^.

The response to SARS-CoV-2 infection involves an interplay of immune pathways that determine disease severity. During the early stages of SARS-Cov2 infection, it is crucial to mount a robust type I interferon response to limit the spread of infection and prime adaptive immune responses that promote viral clearance. In patients with severe disease, delayed and ineffective interferon responses result in increased viral loads. SARS-CoV2 proteins inhibit interferon signalling at multiple steps in the pathway and promote NFkB signalling, resulting in a cytokine storm that drives severe disease^53^. Oxidative stress and excessive tissue damage activates inflammasomes, further driving the cytokine storm, which in turn promotes vascular permeability, decreased oxygen levels in the blood, multi-organ damage, and respiratory failure^53^. *Decipher* analysis of monocyte responses in patients with COVID-19 found that transcription factors that mediate interferon responses (e.g. IRF7, IRF9, STAT1, STAT2) were strongly activated in CD14^+^ monocytes from patients with both mild and severe disease. However, in CD16^+^ monocytes from patients with severe disease, we observed a loss of interferon transcription factors and a gain of NFkB and other factors that mediate responses to oxidative and cellular stress (HIF-1a, ATF3, ATF4). Importantly, this was not observed in patients with mild disease. In this context it is noteworthy that about 10% of blood monocytes are infected with SARS-CoV2 in patients with covid disease^54^. Given that monocytes do not express the viral entry receptor ACE2, virus is taking up by monocytes via antibody-mediated opsonisation through the Fcγ receptor CD16^54^. Once inside monocytes, the viral proteins will inhibit interferon signalling, promote NFkB signalling and inflammasome activation and drive a system cytokine storm.

*Decipher* has limitations that should be acknowledged. We observed intracellular activity to be imbalanced, where some cell types have a greater number of differentially activated TFs. While this is expected behaviour, it may affect the ability to compare prioritization scores between cell types. To address this, a potential alternative could be to perform scaling on TF activity and introduce a normalization factor, in similar manner to scSeqComm^18^. We also observed that TF activity was highly correlated within each cell type. Although multi-collinearity is a known property of biological networks, in the context of our analysis it does pose a problem, as LR pairs that are most predictive of the activity of a TF are likely to be selected for other TFs. Here, we suggest implementing new scoring strategies for TF activity, something that is in active development by the bioinformatics community. Conceptually, *Decipher* currently does not consider the fact that signalling occurs within local niches nor account for feedback control or response dynamics. These issues can be addressed, for example, by extending *Decipher* to spatial transcriptomics data and by modelling this dynamical systems behaviour. Despite these limitations, our findings demonstrate the utility of *Decipher* to decode biological processes and, in doing so, unveil novel therapeutic targets for experimental validation and clinical development. We present *Decipher* as a modular pipeline that quantifies active ligand-receptor pairs, accounts for downstream intracellular responses and maps cell-cell communication at a systems level. We believe that *Decipher* will be invaluable as a tool to accelerate the identification of novel therapeutic targets for human diseases, as well as accelerate the development of new medicines.

## 4 Methods

**Collection and processing of single-cell transcriptomic data of poly-IC stimulation to human blood mononuclear cells from Read et al.**^28^ Processed single-cell data was directly obtained from the original authors. We used the original cell type labels, so no further preprocessing was required. Alternatively, the data are available from Gene Expression Omnibus (Accession Number GSE184383).

**Collection and processing of mouse single-cell transcriptomic data of lung-tissue response to BCG vaccination from Lee et al.**^32^ Count data was downloaded from Gene Expression Omnibus (Accession Number: GSE244126). No cell type labels were provided, so we used ScType^55^ along with two reference tissue profiles (Immune system and lung) to assign cell type labels to each cluster, retaining the label with the largest score for each cluster.

**Collection and processing of human single-cell transcriptomic data of human PBMC response to Pfizer-BioNtech vaccination from Arunachalam et al.**^33^ Count data and phenotypic data were downloaded from Gene Expression Omnibus (Accession Number: GSE171964). We used the original cell type labels, so no further preprocessing was required.

**Collection and processing of single-cell transcriptomic data of PBMCs from subjects with Systemic Lupus (SLE) and healthy control from Perez et al.**^34^ Count data, including phenotypic data, was downloaded from Gene Expression Omnibus (Accession number: GSE174188). We used the original cell type labels, so no further preprocessing was required. We constrained our analysis to the subset of samples belonging to female individuals of Asian ancestry that were either classified as healthy or as ‘managed’ SLE cases.

**Collection and processing of single-cell transcriptomic data of PBMCS from ICU patients with and without sepsis from Reyes et al.**^29^ Count data with phenotypic information were downloaded from the Broad Institute Single-cell Portal (Accession Number: SCP548). We used the original cell type labels, so no further preprocessing was required.

**Collection and processing of single-cell transcriptomic data of PBMCS from breast cancer patients treated with immune checkpoint blockade from Bassez et al.**^38^ Processed single-cell data was downloaded directly from Diether Lambrecht’s laboratory website, through their Data portal. We used the original cell type labels, so no further preprocessing was required.

**Collection and processing of single-cell transcriptomic data of PBMCS from patients with mild or severe COVID-19 from Arunachalam et al**.^31^ Count data of PBMCs from mild and severe COVID-19 patients was downloaded from GEO (accession number: GSE155673). Only samples corresponding to actual RNA-seq experiments were selected (cov01–cov04, cov07–cov12, cov17–cov18). For each sample, the corresponding barcodes.tsv.gz and matrix.mtx.gz files were retrieved. Separate Seurat objects were created for each sample and merged into a single object, with sample identifiers tracked in metadata. Sample-level metadata (age, sex, disease status, severity, and days since symptom onset) were curated from GEO and merged with cell-level metadata. We followed the preprocessing protocol outlined in the original study to produce cluster-level marker genes distinguishing severe and mild COVID-19 from Healthy controls. We applied the Azimuth pipeline^47^ to label cell types (at level-2 resolution).

**Collection and processing of single-cell transcriptomic data from the CellxGene collection**^56^. Processed single-cell datasets were downloaded as annotated .h5ad objects from the CZ CellxGene portal. For each dataset, we applied consistent preprocessing steps: cells with fewer than 200 detected genes were removed. Datasets were optionally subset based on disease or condition annotations, depending on the study context. The AnnData objects were parsed in Python, and we verified that raw count matrices contained integer values. Metadata, gene names, and filtered count matrices were extracted and saved in standard formats. Where applicable, Ensembl gene IDs were converted to HGNC symbols. Resulting count matrices and metadata was used to create Seurat objects in R, with cleaned cluster annotations and standardized condition labels. Original cell type and disease labels were retained throughout

This workflow was applied to: PBMCs from healthy individuals and patients with influenza from Lee et al.^30^; Pancreatic islets from healthy donors and individuals with type 1 diabetes from Fasolino et al.^35^ ; Head and neck squamous cell carcinoma samples from Jenkins et al.^39^; Kidney samples from healthy individuals and chronic kidney disease patients from Lake et al.^37^; and Bronchial biopsy samples from cystic fibrosis patients and healthy controls from Berg et al.^36^.

### Preprocessing pipeline

If needed, a standard preprocessing pipeline based on Seurat^47^ is delineated in “Supplementary Note 1: Preprocessing pipeline for single-cell profiles”. Said pipeline addresses standard scRNA-seq preprocessing, beginning after the raw data has been read aligned to produce gene counts ***D***. In brief, this preprocessing pipeline covers the filtering out poor-quality, invalid, or damaged cells, normalization of gene counts, using these counts to produce clusters 𝔾_*j*_ of similar cells and determining representative genes for each cluster for downstream cell-labelling. To aid with interpretability of results, we recommend clusters are labelled using expert judgement. If expert judgement is not available, there are several cell type labelling tools, one of which we used (ScType^55^) to pre-process lung-tissue data from Lee et al^32^.

### Meta-cell generation

To remove noise and account for zero-inflation of gene counts in scRNA-seq data, we applied the method proposed by Baran et al.^25^ with parameters determined from guidelines by Obradovic et al.^57^ Here, randomly-sampled individual single-cell profiles from each condition and cell type were aggregated into meta cells by adding the counts of each gene across similar cells (based on Euclidean distance of their gene counts). The number of cells per meta-cell, *k*, was determined such that for most cell types, the median gene count per cell was between 7,500 and 10,000^57^. We constrained the number of meta-cells such that each cell type had the same number of meta-cells, and, on average, no cell appeared in more than one meta-cell. In addition, we include a parameter to limit the number of meta-cells generated for each cluster-condition pair. This prevents overrepresentation of large cell groups and ensures computational tractability by capping the number of meta-cells at 600 per group, which typically exceeds the number of features modelled in downstream analyses. Lastly, each meta-cell was normalized by its new library size and scaled by a factor of 10^’^for interpretability.

### Interaction potential between clusters

As a pre-requisite to determine which ligand-receptor (LR) pairs are actively modulating a system’s response, we first confirmed that these same LR pairs had the potential to signal. Here, we relied on the reference LR pair database ConnectomeDB2020^22^. LR interactions in ConnectomeDB2020 were manually curated, based on literature support, and focused on monomeric interactions, i.e. a single ligand interacts with a single receptor. This aligned with our current version of the pipeline, as *Decipher* currently does not support multimeric interactions.

We retained any ligand that was expressed in at least 10% of cells in any cell type and condition. We addressed ligands at a global level because we assumed that ligand-concentration in the environment is the driver of signalling, as opposed to ligand production by a single cluster. Receptors were filtered on a cluster-by-cluster basis. For each cell type, a receptor was only retained if it was expressed in at least 10% of the cells in either condition. Receptors were treated as cluster-specific because receptor signalling is biologically constrained to individual cells. Together, this yielded a cluster-specific list of LR pairs.

The condition-specific interaction potential *p* of a ligand-receptor pair *i* in each meta-cell from a receiver cell type was calculated as:

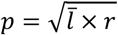

Where *l̅* represents the mean ligand expression of ligand *l* across all cell types in each condition and *r* represents the receptor expression in a meta-cell. We performed the Wilcoxon Rank Sum test as implemented in the *FindMarkers* function from the *Seurat* package to identify differentially expressed interactions (*p*_*adj*_ < 0.01 & log_2_(⋅) > 0.1) across conditions. Here, we assumed that the interaction potential had to exhibit some change across conditions to be relevant for downstream analysis.

A significant portion of selected LR pairs exhibited high correlation with each other and thus could affect downstream statistical modelling. Highly correlated LR pairs occurred primarily when multiple ligands shared a single receptor (Supplementary Fig. SF.2). Therefore, we focused on selecting representative interactions for those interactions whose receptors had multiple complementary ligands. To identify which interactions behaved similarly across samples, we calculated the distance between each pair of interactions as:

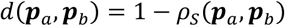

Where ***p***_*a*_ and ***p***_*b*_ represent the vector of interactions potentials for two LR pairs, and *ρ*_*s*_(⋅) represents the Spearman correlation function. We performed hierarchical clustering on these distances to define clusters of similarly behaving interactions. Subsets produced by hierarchical clustering were defined using a distance metric threshold; we set this parameter so that we produced approximately the same number of clusters as unique receptors within our features. Lastly, we randomly selected and retained one representative LR interaction from each subset.

### Context-specific GRNs for each cluster

We relied on base gene regulatory networks provided by *CellOracle*^23^, as these were generated using single-cell ATAC-seq data which, unlike computationally derived gene regulatory networks, ensures there is biological evidence of a relationship between a transcription factor (TF) and a target gene (TG). We further relied on *CellOracle* to tailor this network to each cell type. *CellOracle* is implemented as a Python pipeline, which required converting our Seurat objects to Python objects. Given difficulties with mapping Seurat objects to the required format, we converted our count matrices to Python-compatible matrices and re-processed these counts using the Scanpy^58^ pipeline. The pipeline consisted of removing genes with no counts, normalizing cell counts by library size, filtering normalized counts to retain only the top 3,000 most variable genes, which were re-normalized by their new library size, and taking the natural logarithm and scaling these counts to unit variance and zero mean. We ran the *get_links()* function from the *cellOracle* pipeline on the scaled counts to tailor the base gene regulatory network to each cell type.

The tailored gene regulatory networks produced by *CellOracle* proved too dense, so we only retained the strongest 20,000 edges in the network (based on the absolute values of their weights). This first pruning was a more global, entire-network based-pruning. We complemented this global pruning with a more local pruning of edges, such that for each TF present in the network, we restricted its outdegree to a maximum of 40 edges, retaining the largest edges within each sub-network. We performed pruning of the network to increase the heterogeneity of regulons (as large regulons naturally tended to have overlap with smaller regulons.

### Scoring TF activity

Using the tailored networks, we proceeded to calculate the activity of individual TFs based upon the expressions of their TGs using elements of PAGODA2^27^. We applied PAGODA2’s ‘mixture model’ to the meta-cell count matrix to derive a matrix of residuals, capturing the over- or under-dispersion of each gene in each meta-cell. We applied PAGODA2’s function *testPathwayOverdispersion()* to the residual matrix. This function is an implementation of the implicitly restarted Lanczos bidiagonalization algorithm, which returns the first principal component for a given. For each meta-cell, we projected its residual vector onto that first principal component, obtaining a meta-cell-specific activity score. The collection of these scores across all meta-cells was then standardized to yield a TF activity score.

Relevant TFs for each cell-type were then selected based on statistically significant deviations in their activity across conditions. Specifically, we assessed significance (two-tailed, *p* < 0.05) by comparing TF activity scores to a null distribution of scores generated from randomized gene regulatory networks, in which target genes were shuffled independently for each TF while preserving the number of targets.

### Building integrated signalling networks

For each cluster, we began by assuming that LR pairs and TFs formed a directed bipartite network where each ligand-receptor pair had the potential to influence every transcription factor. We then refined this network by inferring both the weights and mode of influence (direction) of each edge. To do so, we modelled the relationship between LR pairs, represented by their interaction potentials, and transcription factors, represented by their TF activity profiles.

For each cell type, we formulated this as *n* separate regression problems, one for each TF *f*, represented by its TF activity vector, ***t***_*f*_. For each TF *f*, we modelled its relationship with all LR pairs signalling potential ***P*** relevant to that cell type as:

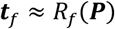

Where *R_f_*(⋅) represents the regression function mapping ligand-receptor interaction potentials to the activity of TF *f*. Here, we used a Random Forest regressor.

For each regression problem, we further calculated each feature’s permutation importance metric as:

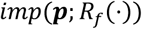

Where *imp*(⋅) represents the importance function, which randomly shuffles the interaction potential vector ***p*** to calculate the impact of permuting a given interaction on model performance. The importance metric was used as the weight of the edge between LR pairs and transcription factors. Furthermore, as LR pairs can either activate or suppress TF activity, we estimated the mode of influence for each edge as the sign of the Spearman correlation between each predictor-target pair in the model.

Repeated across *n* problems this yields a weighted adjacency matrix ***A*** connecting all LR interactions and TFs in each cell type. Naturally, it is possible to extend this network to include target genes, as this information is already encoded in the tailored, cluster-specific gene regulatory networks.

### Scoring differential ligand-receptor activity

Having obtained cluster-specific signed, weighted, and directed iSNs comprising ligands, receptors and TFs, we utilized these networks to score LR interactions for each cell type. Since *Decipher* was designed to compare signalling at the cell type level, we first calculated the changes in intracellular TF activity for each cluster as the difference in median TF activity for each TF *f* across conditions, Δ*_f_*.

Differential activity is then captured by the *Decipher* score S, which is represented by:

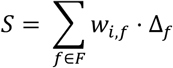

Where *F* represents all TFs relevant to a given cell type, and *w*_*i,f*_ represents the weight between a ligand-receptor pair *i* and a TF *f*.

Taken altogether, with this approach, we achieve our two primary aims of building cluster-specific Integrated Signalling Networks and scoring interaction activity based on such a network.

### CytoSig ligand activity

We estimated ligand activity with CytoSig^42^. For each cluster, we only retained genes with non-zero total counts across all cells. Gene counts were normalized by total cell count, scaled by a factor of 100,000, and log-transformed using log_2_(*x* + 1). We defined a differential expression profile for each case cell by subtracting the mean expression of control cells from the log-transformed expression values. We applied CytoSig to the differential expression profiles of cells in each cluster, producing z-scores for the ligand activity in each cell.

### Evaluation of ligand-activity prediction

We evaluated how well ligand-receptor predictions from each method aligned with the CytoSig model. Here, ligands were assumed to be active if they displayed an absolute median z-score > 2. Most methods reported scores for some ligands but not others, so we assumed that no prediction implied no activity, and assigned these ligands zero values. Except *Decipher*, all methods output scores for each sender-receiver cell type pair. Therefore, we reduced these scores to a single score per ligand and receiver cell type by selecting the LR interaction with the highest absolute score involving that ligand.

## Acknowledgements

We would like to acknowledge Dr. J Heng from Remotely Consulting for the services provided in English-language editing of the manuscript, as well as for his valuable feedback on *Decipher*.

## Notes

### Competing Interest Statement

AB is the founder of the start-up company INSiGENe Pty Ltd that funded this work. AB is a co-founder, equity holder, and director of the startup company Respiradigm Pty Ltd that is unrelated to this work. EB received a scholarship from INSiGENe Pty Ltd to conduct this work.

### Summary of Updates

This version of the manuscript incorporates multiple changes. Additional datasets have been included in the benchmarking, providing a broader evaluation of method performance. We also include the in-detail analysis of a study profiling mild and severe COVID-19. Several sections from the earlier version have been removed to improve focus, including the comparison with a broader set of cell-cell communication tools and the evaluation of context-specific integrated networks.

