## Supplementary information for "Decipher: A computational pipeline to extract context-specific mechanistic insights from single-cell profiles"

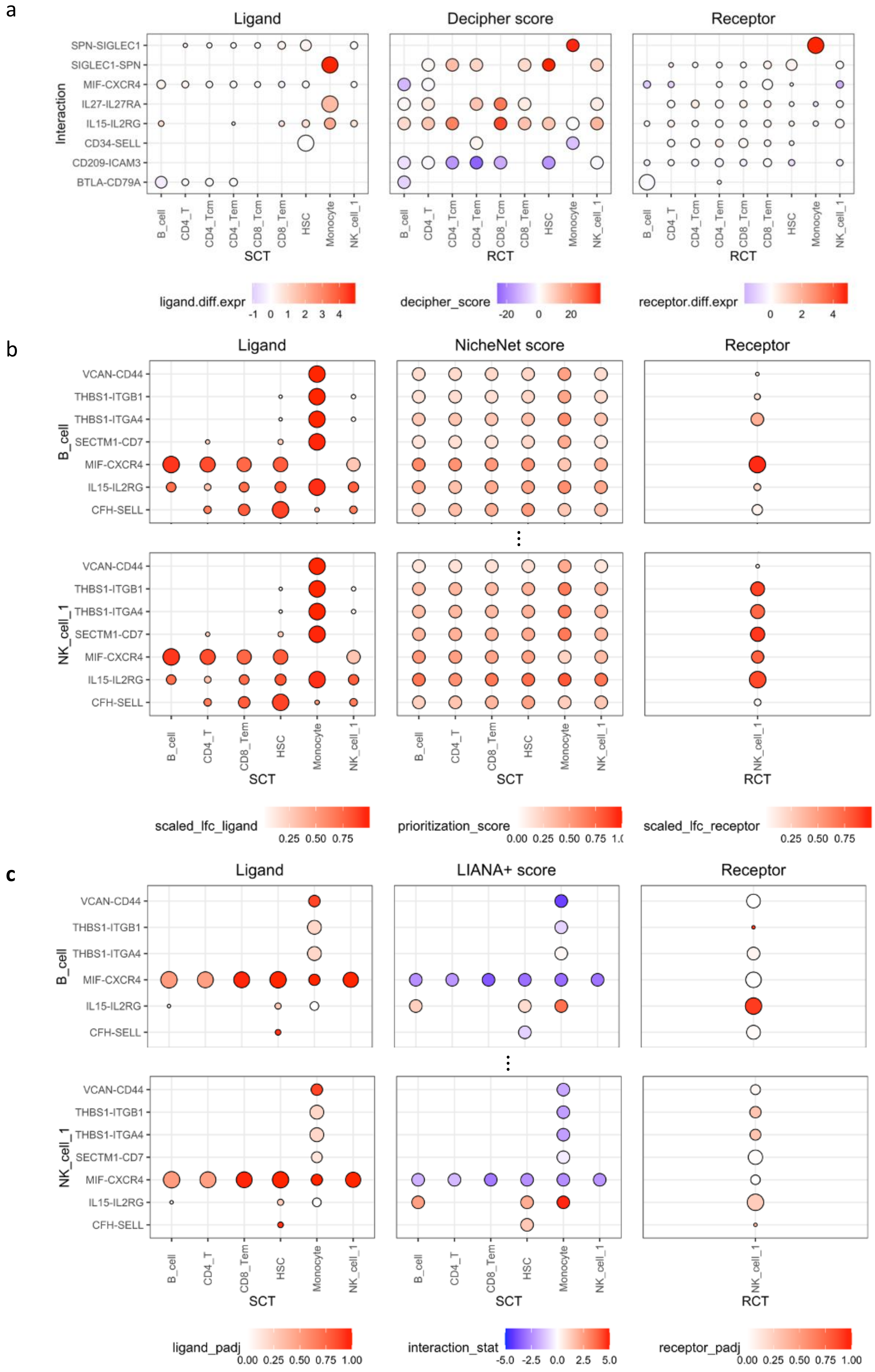

a
Ligand
Decipher score
Receptor
Interaction
SPN-SIGLEC1
SIGLEC1-SPN
MIF-CXCR4
IL27-IL27RA
IL15-IL2RG
CD34-SELL
CD209-ICAM3
BTLA-CD79A
B_cell
CD4_T
CD4_Tcm
CD4_Tem
CD8_Tcm
CD8_Tem
HSC
Monocyte
NK_cell_1
SCT
RCT
ligand.diff.expr
-1 0 1 2 3 4
decipher_score
-20 0 20
receptor.diff.expr
0 2 4
b
Ligand
NicheNet score
Receptor
B_cell
NK_cell_1
VCAN-CD44
THBS1-ITGB1
THBS1-ITGA4
SECTM1-CD7
MIF-CXCR4
IL15-IL2RG
CFH-SELL
B_cell
CD4_T
CD8_Tem
HSC
Monocyte
NK_cell_1
SCT
RCT
scaled_lfc_ligand
0.25 0.50 0.75
prioritization_score
0.00 0.25 0.50 0.75 1.0
scaled_lfc_receptor
0.25 0.50 0.75
c
Ligand
LIANA+ score
Receptor
ligand_padj
0.00 0.25 0.50 0.75 1.00
interaction_stat
-5.0 -2.5 0.0 2.5 5.0
receptor_padj
0.00 0.25 0.50 0.75 1.00

**Supplementary Figure 1 (SF.1).**

Comparison of systems-level CCC maps for Cord PIC versus Unstimulated Cord CBMC. Our 'case' mimics a viral infection in Cord Monocytes cells, whereas our 'control' represents Cord Monocytes in their natural state. Systems-level map for all SCTs and RCTs of the top eight (four positive, four negative) LR interactions as determined by Decipher. Results from (a) *Decipher*, (b) NicheNet[1] and (c) LIANA+[2] in Decipher's visualization format, as shown. The format is based on three sub-plots, the LHS sub-plot displays ligand-level statistics for the SCT and the RHS sub-plot presents receptor statistics for the RCT. We utilized the statistics suggested by each method to colour and size the bubbles in these plots. The central sub-plot presents each method's prioritization scores for each ligand in either (a) each RCT (b,c) each SCT-RCT pair. NicheNet and LIANA+ results for each RCT are displayed in a separate row indicated by the LHS label of each plot.

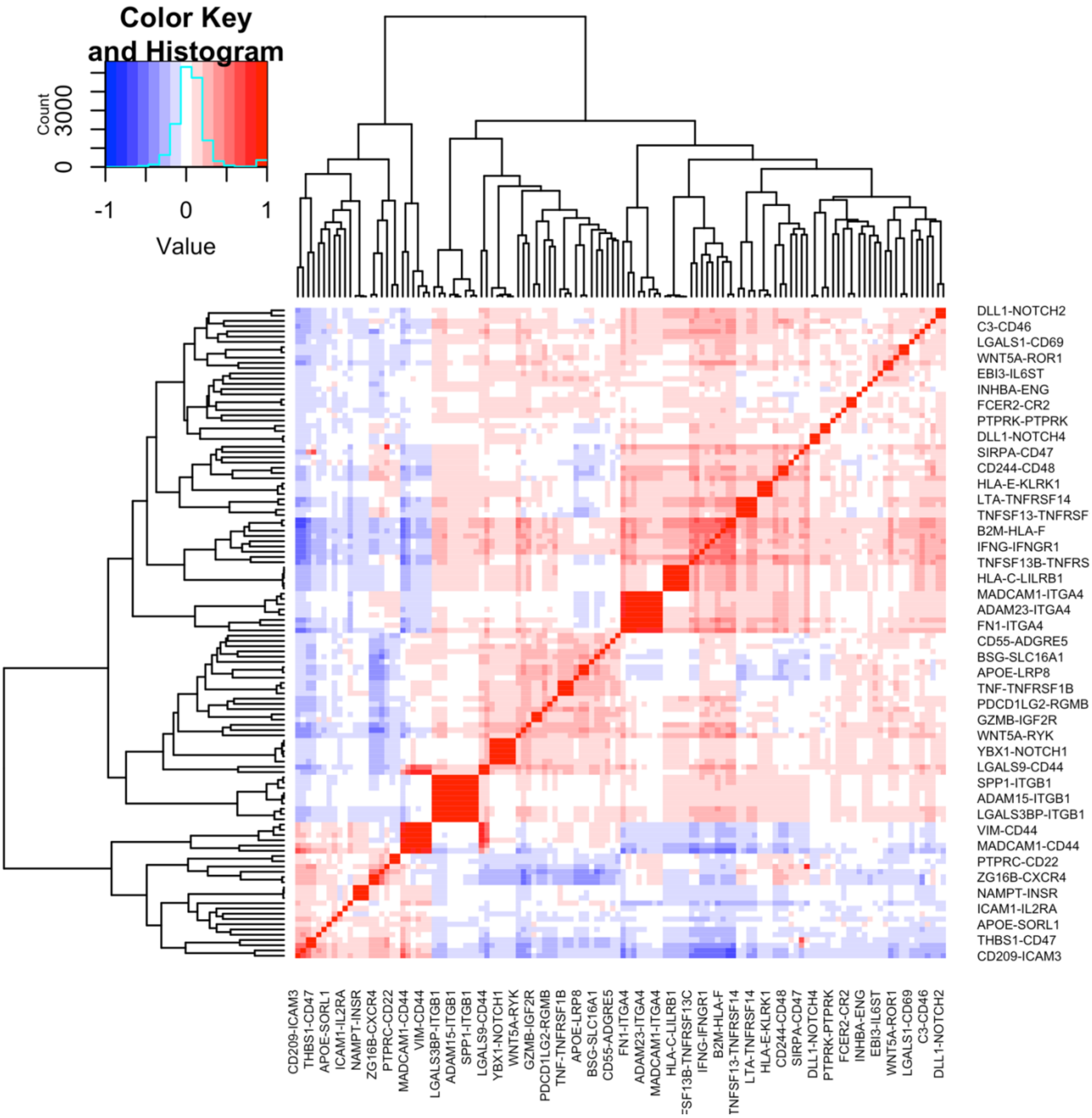


**Supplementary Figure 2 (SF.2).**

Correlation of interaction potential LR pairs. Heatmap of correlation among LR pairs for the B cell cluster in the poly-IC treated CBMC dataset, illustrating that the receptor in clusters of high correlation tends to coincide. For example, {VIM-CD44, MADCAM1-CD44}, {SPP1-ITGB1, ADAM15-ITGB1, LGALS3BP-ITGB1}.

### Supplementary Note 1: Pre-processing pipeline for single-cell profiles

Here we outline a standard implementation of the pre-processing pipeline proposed by Seurat[3], formalizing the steps so as to be compatible with the *Decipher* pipeline.

**Input single-cell transcriptomic expression data.** We define input data as the RNA seq count matrix $\boldsymbol{D}$, where each column $\boldsymbol{d}_c = \left[d_1, \cdots, d_g\right]^T$ presents the gene counts $d_g$ of each gene $g$ in the $c^{th}$ cell. Additional information is represented as an associated list of traits which indicate either the physiological components of the cell itself or the subject from whom the cell is derived from. For instance, if the cell comes from a diseased subject, traits may include age, gender, type of disease, and disease severity. Whereas cell-specific traits may indicate that cell's type and differentiation stage.

**Filtering gene counts.** Filtering is based on general heuristics and is designed to filter out poor quality cells. Here, cells are filtered out if they match any of the following three criteria: (i) present non-zero counts for more than 2,500 genes, indicating that the observed cell probably accounts for more than one cell. (ii) Exhibit non-zero counts for less than 200 genes, which suggests that the cell may be either be damaged or $\boldsymbol{D}_c$ is background noise. (iii) Displays greater than 5% of their total counts to be coming from mitochondrial genes, implying poor sample quality.

**Normalizing gene counts.** Normalization, scaling and log-transformation of the count matrix is done using the following eq.

$$\overline{\boldsymbol{d}}_c = \log(\boldsymbol{d}_c) - \log(\alpha_c) + \log\,(10^6)$$

Where $\alpha$ represents the library size of the cell (the total RNA content), $10^6$ is a scaling factor to aid with interpretability of normalized counts and the logarithmic transformation aids with (i) stabilizing variance across different levels of expression (ii) making the data more normally distributed, and (iii) reduces the impact of extreme values. Applied to each cell, this yields the normalized count matrix $\overline{\boldsymbol{D}}$.

**Defining similarity among cells.** To cluster cells into similar groups, we first define similarity among cells. Here, we rely on principal component analysis (PCA) on the top variable features of $\overline{\boldsymbol{D}}$ (2000 by default), from which we retain the top 30 principal components, $\overline{\overline{\boldsymbol{D}}}$. This reduced matrix is used to construct a distance matrix $\boldsymbol{E}$, representing the pair-wise Euclidean distances measured for all cells with respect to one another. From $\boldsymbol{E}$, we identify the top $k$ nearest neighbours for each cell $C_a$, $\mathcal{S}_a = \{C_{a,1}, C_{a,2}, \cdots, C_{a,k}\}$ (20 by default). The neighbourhood of each cell is used to measure the similarity of any two cells Jaccard Similarity index. Given two cells $C_i$ and $C_j$, with their respective neighbourhoods i.e., $\mathcal{S}_i$ and $\mathcal{S}_j$, their respective overlap $n_{i,j}$ and differences $n_i$, $n_j$ are used to evaluate the following index

$$\mathcal{J}_{i,j} = \frac{n_{i,j}}{n_i + n_{i,j} + n_j}$$

where said index is meant to represent the similarity of two cells. By default, the matrix $\mathcal{J}$ of Jaccard similarity indices is pruned such that any $\mathcal{J}_{i,j} \leq \frac{1}{15}$ is reduced to 0. For convenience, $\mathcal{J}$ can be thought of as a graph connecting cells with edges whose weight represents their similarity.

**Clustering cells based on $\mathcal{J}$.** To cluster cells into communities presenting similar traits, Seurat leverages the Louvain algorithm. The algorithm adopts a greedy approach wherein it optimizes a '*modularity index*' ($Q$) that measures the density of edges within clusters in relation to the density of edges between clusters. Here, the modularity index for a cluster $\mathbb{G}_j$ (a subset of cells) is taken to be

$$Q_{\mathbb{G}_j} = \frac{A}{2 \cdot m} - \left(\frac{B}{2 \cdot m}\right)^2$$

Where $m$ is the sum of all edges in the graph $\mathcal{J}$, $A$ is the sum of all edges in $\mathcal{J}$ connecting cells only within the cluster $\mathbb{G}_j$, and $B$ is the sum of all elements in $\mathcal{J}$ connecting any cell to cells in $\mathbb{G}_j$.

Similarly, the change in modularity index ($\Delta Q$) represents the difference in $Q$ when the $i^{th}$ cell $C_i$ is removed from cluster $\mathbb{G}_j$ and assigned to another cluster $\mathbb{G}_k$.

The algorithm works by assigning each cell to its own cluster then, for each cell, iteratively assigning that cell to its neighbour's clusters and calculating the respective $\Delta Q$. If the change in modularity for all assignments is negative, then the cell is kept in its original cluster, otherwise it is assigned to the cluster that leads to the largest change in modularity. We refer to the final clustering by $\mathbb{G}_j$.

**Determine representative genes for each cluster.** For each cluster $\mathbb{G}_j$, Seurat employs the Mann-Whitney U-statistic to determine the set of genes that best represent that cluster. Where the U-statistic is used to calculate a p-value on the observed value of a gene for a particular cluster which, along with the $\log_2(\cdot)$ fold change, is used to determine representative genes $\mathcal{S}_{\mathbb{G}_j}$ of cluster $\mathbb{G}_j$.

**Assign attributes to each cluster using expert judgement:** $\mathcal{S}_{\mathbb{G}_j}$ is utilized by expert judgement to assign attributes that best denote each $\mathbb{G}_j$. In this case, such attributes primarily correspond to the biological cell-type that best fits the cluster.

1. Browaeys, R., Saelens, W. & Saeys, Y. NicheNet: modeling intercellular communication by linking ligands to target genes. *Nat Methods* **17**, 159–162 (2020).
2. Dimitrov, D. *et al.* LIANA+ provides an all-in-one framework for cell–cell communication inference. *Nat Cell Biol* **26**, 1613–1622 (2024).
3. Hao, Y. *et al.* Integrated analysis of multimodal single-cell data. *Cell* **184**, 3573-3587.e29 (2021).
